# TESS: Bayesian inference of lineage diversification rates from (incompletely sampled) molecular phylogenies in R

**DOI:** 10.1101/021238

**Authors:** Sebastian Höhna, Michael R. May, Brian R. Moore

**Affiliations:** Department of Evolution and Ecology, University of California, Davis, CA 95616, USA; Department of Integrative Biology, University of California, Berkeley, CA 94720, USA; Department of Statistics, University of California, Berkeley, CA 94720, USA; Department of Mathematics, Stockholm University, Stockholm, SE-106 91, Sweden

## Abstract

**Summary:** Many fundamental questions in evolutionary biology entail estimating rates of lineage diversification (speciation – extinction). We develop a flexible Bayesian framework for specifying an effectively infinite array of diversification models—where rates are constant, vary continuously, or change episodically through time—and implement numerical methods to estimate parameters of these models from molecular phylogenies, even when species sampling is incomplete. Additionally we provide robust methods for comparing the relative and absolute fit of competing branching-process models to a given tree, thereby providing rigorous tests of biological hypotheses regarding patterns and processes of lineage diversification.

**Availability and implementation:** the source code for TESS is freely available at http://cran.r-project.org/web/packages/TESS/.

## 1 Introduction

Stochastic-branching process models (*e.g.,* birth-death models) describe the process of diversification that gave rise to a given study tree, and include parameters such as the rate of speciation and extinction. Parameters of these models are commonly estimated from molecular phylogenies using maximum-likelihood methods (*e.g.*, Paradis *et al.*, 2004; Rabosky, 2006; Stadler, 2013). There are several potential benefits of pursuing this inference problem in Bayesian statistical framework, such as: (1) providing a natural means for accommodating uncertainty in our estimates (by inferring parameters as posterior probability densities rather than point values); (2) incorporating prior information regarding various aspects of the branching-process models (such as the expected number or severity of mass-extinction events), and; (3) leveraging robust Bayesian approaches for model comparison and model averaging.

These considerations influenced our development of TESS, an R package for the Bayesian inference of lineage diversification rates that allows researchers to address three fundamental questions: (1) *What are the rates of the process that gave rise to my study tree?* (2) *Have diversification rates changed through time in my study tree?* (3) *Is there evidence that my study tree experienced mass extinction?*

## 2 Methods and algorithms

### Branching-process models

Inferring rates of lineage diversification is based on the *reconstructed evolutionary process* described by Nee *et al.* (1994); a birth-death process in which only sampled, extant lineages are observed. Our implementation exploits recent theoretical work (Lambert, 2010; Höhna, 2013, 2014, 2015) that allows the rate of diversification to be specified as an arbitrary function of time. By virtue of adopting this generic approach, it is possible to specify an effectively infinite number of branching-process models in TESS. These possibilities correspond to four main types of diversification models: (1) constant-rate birth-death models; (2) continuously variable-rate birth-death models; (3) episodically variable-rate birth-death models, and; (4) explicit mass-extinction birth-death models.

### Phylogenetic data

Parameters of the branching-process models are inferred from a given study tree. Specifically, TESS takes as input rooted *ultrametric* trees, where all of the tips are sampled at the same time horizon (the present). Other types of trees—*e.g.,* where tips are sampled sequentially through time (Heath *et al.*, 2014)—are currently not supported. It is now well established that estimates of diversification rates are sensitive to incomplete species sampling (*i.e.,* where the study tree includes only a fraction of the described species; Cusimano and Renner, 2010; Höhna *et al.*, 2011). This is a particular concern, as most empirical phylogenies include only a fraction of the member species. Accordingly, TESS implements various approaches for accommodating incompletely sampled trees, including uniform sampling and diversified sampling schemes (Höhna *et al.*, 2011; Höhna, 2014).

### Parameter estimation

In TESS, parameters of the branching-process models are inferred in a Bayesian statistical framework. Specifically, we estimate the joint posterior probability density of the model parameters from the study tree using numerical methods—Markov chain Monte Carlo (MCMC) algorithms (Figure 1). The numerical methods implemented in TESS include adaptive-MCMC algorithms (Haario *et al.*, 1999)—where the scale of the proposal mechanisms is automatically tuned to ensure optimal efficiency (mixing) of the MCMC simulation—and also feature real-time diagnostics to assess convergence of the MCMC simulation to the stationary distribution (the joint posterior probability density of the model parameters).

**Fig. 1.**
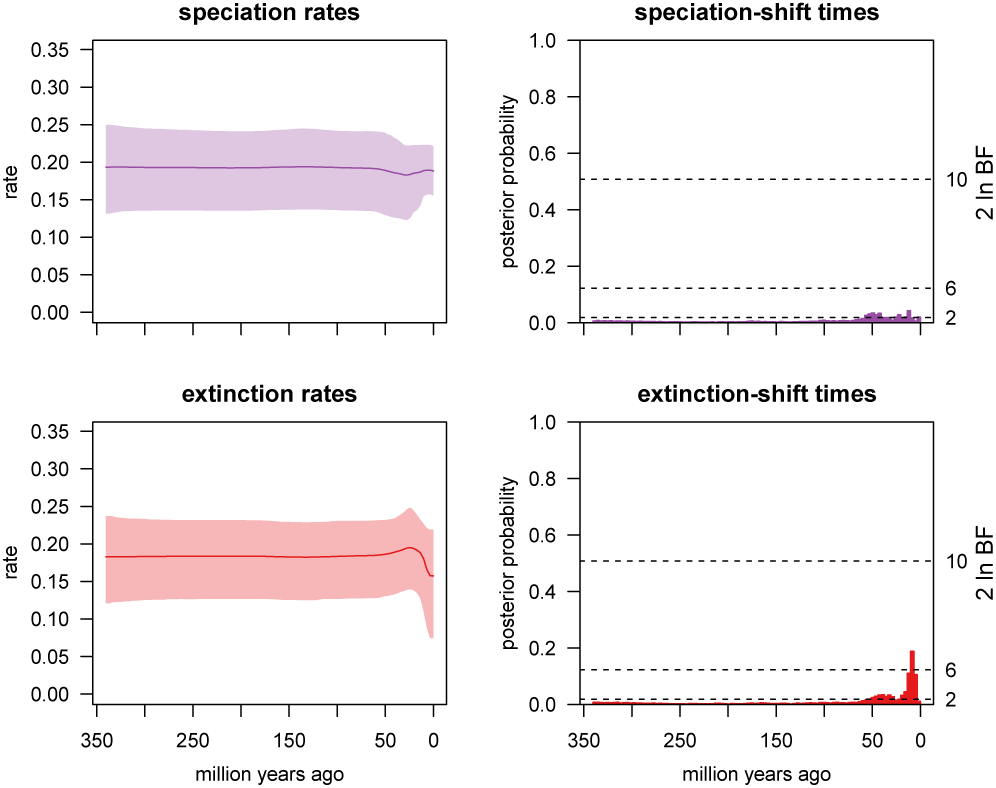
Estimating rates of (and identifying shifts in) lineage diversification through time. Left: Plots of the posterior mean and 95% credible interval for the speciation and extinction rate (upper and lower panels, respectively). Right: Identifying temporal shifts the speciation and extinction rate (upper and lower panels, respectively). Each bar indicates the posterior probability of at least one rate shift within that interval. Bars that exceed the specified significance threshold (here, 2 ln BF *>* 6) indicate significant rate shifts. This analysis of the conifer tree from Leslie *et al.* (2012) reveals a significant shift in the extinction rate *∼* 5 million years ago.

### Model comparison

Each branching-process model specifies a possible scenario for the diversification process that gave rise to a given study tree. For most studies, several (possibly many) competing branching-process models of varying complexity will be plausible *a priori*. We therefore need a way to objectively identify the best candidate diversification model. Bayesian model selection is based on *Bayes factors* (*e.g.,* Kass and Raftery, 1995; Suchard *et al.*, 2001; Holder and Lewis, 2003). This procedure requires that we first estimate the marginal likelihood of each candidate model, and then compare the ratio of the marginal likelihoods for each pair of candidate models. We have implemented both *stepping-stone sampling* (Xie *et al.*, 2011; Fan *et al.*, 2011) and *path-sampling* (Lartillot and Philippe, 2006; Baele *et al.*, 2012) algorithms for estimating the marginal likelihoods of branching-process models in TESS, which provides a robust and flexible framework for Bayesian tests of diversification-rate hypotheses.

### Model adequacy

Bayes factors allow us to assess the *relative* fit of two or more competing branching-process models to a given study tree. However, even the very best of the competing models may nevertheless be woefully inadequate in an *absolute* sense. Accordingly, TESS implements methods to assess the absolute fit of a candidate diversification model to a given study tree using *posterior-predictive simulation* (Gelman *et al.*, 1996; Bollback, 2002; Moore and Donoghue, 2009; Brown, 2014). The basic premise of this approach is as follows: if the diversification model under consideration provides an adequate description of the process that gave rise to our study tree, then we should be able to use that model to generate new phylogenies that are in some sense ‘similar’ to our study tree. TESS permits use of any summary statistic—*e.g.,* the *γ*-statistic (Pybus and Harvey, 2000) or the nLTT statistic (Janzen *et al.*, 2015)—to measure the similarity between predicted and observed data.

### Model averaging

The vast space of possible branching-process models precludes their exhaustive pairwise comparison using Bayes factors. This issue may be addressed by means of model-averaging approaches that treat the model as a random variable (Huelsenbeck *et al.*, 2004, 2006). TESS implements such an approach; the CoMET (CPP on Mass-Extinction Times) model (May *et al.*, 2015). The CoMET model is comprised of three compound Poisson process (CPP) models that describe three corresponding types of events: (1) instantaneous tree-wide shifts in speciation rate; (2) instantaneous tree-wide shifts in extinction rate, and; (3) instantaneous tree-wide mass-extinction events. The dimensions of the CoMET model are therefore dynamic; there is effectively an infinite number of nested models that include zero or more events. We use reversible-jump MCMC to average over all possible models, visiting each model in proportion to its posterior probability (Green, 1995; Huelsenbeck *et al.*, 2000). The resulting joint posterior probability distribution can then be queried to assess whether the study tree has been impacted by mass extinction, and if so, to identify the number and timing of those events using Bayes factors (Figure 2).

**Fig. 2.**
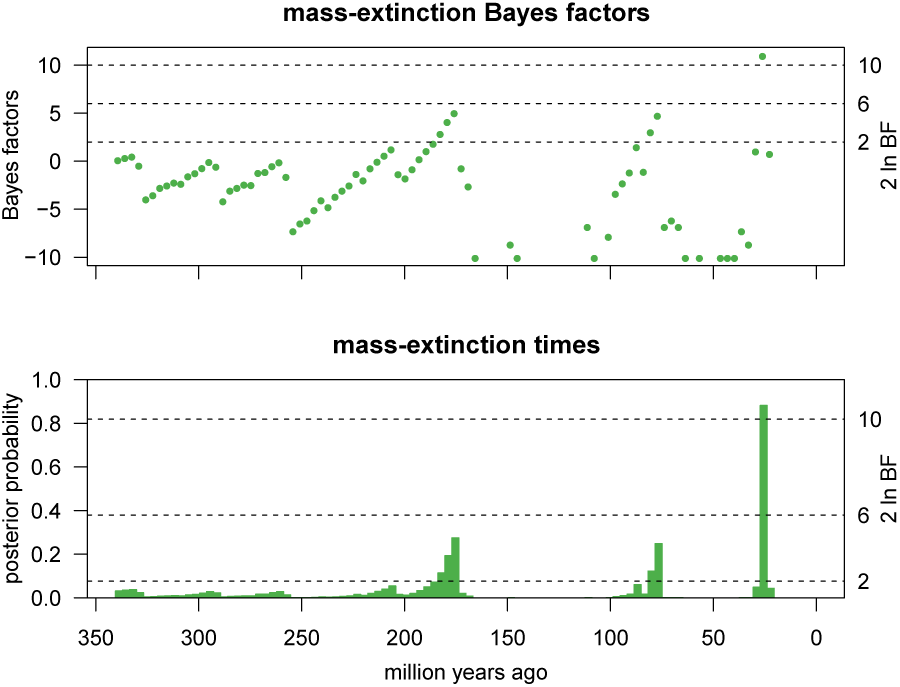
Identifying significant mass-extinction events using Bayes factors. Each bar indicates the posterior probability of at least one mass extinction within that interval. Bars that exceed the specified significance threshold (here, 2 ln BF *>* 6) are inferred to be significant mass-extinction events. This analysis of the conifer tree from Leslie *et al.* (2012) identifies two significant mass-extinction events that occurred 48 and 93 million years ago.

## 3 Conclusions

TESS allows users to specify an effectively countless number of diversification models, where each model describes an alternative scenario for the diversification of the study tree. Additionally, TESS provides robust methods for assessing the relative fit of competing models to a given study tree, providing users with an extremely flexible yet intuitive framework for testing hypotheses regarding the patterns and processes of lineage diversification. We are optimistic that the implementation of a robust and powerful Bayesian statistical framework for exploring rates of lineage diversification will provide biologists with an important tool for advancing our understanding of the processes that have shaped the Tree of Life.

## Acknowledgements

*Funding*: This research was made possible by NSF grants DEB-0842181, DEB-0919529, DBI-1356737, and DEB-1457835 awarded to BRM, and by a Miller Institute for Basic Research in Science scholarship awarded to SH.

*Conflict of interest*: None declared.

